# Non-invasive vagus nerve stimulation and the motivation to work for rewards: a replication

**DOI:** 10.1101/2023.03.13.532362

**Authors:** Federica Lucchi, Beth Lloyd, Sander Nieuwenhuis

## Abstract

**Background:** The vagus nerve is thought to be involved in the allostatic regulation of motivation and energy metabolism via gut-brain interactions. A recent study by Neuser and colleagues [1] provided novel evidence for this process in humans, by reporting a positive effect of transcutaneous auricular vagus nerve stimulation (taVNS) on the invigoration of reward-seeking behaviors, especially for food rewards.

**Objective:** We conducted an independent direct replication of Neuser et al. [1], to assess the robustness of their findings.

**Methods:** Following the original study, we used a single-blind, sham-controlled, randomized cross-over design. We applied left-sided taVNS in healthy human volunteers (n=40), while they performed an effort allocation task in which they had to work for monetary and food rewards. The replication study was purely confirmatory in that it strictly followed the analysis plans and scripts used by Neuser et al. [1].

**Results:** Although, in line with Neuser et al. [1], we found strong effects of task variables on effort invigoration and effort maintenance, we failed to replicate their key finding: taVNS did not increase the strength of invigoration (*p* = .62); the data were five times more likely (BF_10_ = 0.19) under the null hypothesis. We also found substantial evidence against an effect of taVNS on effort maintenance (*p* = 0.50; BF_10_ = 0.20).

**Conclusions:** Our results provide evidence against the idea that taVNS boosts the motivational drive to work for rewards. Our study also highlights the need for direct replications of influential taVNS studies.

## Introduction

Transcutaneous auricular vagal nerve stimulation (taVNS) is a non-invasive technique that electrically stimulates the auricular branch of the vagus nerve, the nerve that connects the gut and brain through the gut-brain axis. The stimulation is applied to the outer ear, at the cymba concha, on the inner side of the tragus. Depending on stimulation parameters, taVNS has been found to activate the nucleus of the solitary tract (NTS) and brainstem arousal nuclei via afferent vagal pathways [2,3]. taVNS is used as a treatment of drug-resistant epilepsy and depression [4,5], and has promising effects on other clinical outcome measures [6,7].

Laboratory experiments have also reported positive effects of taVNS on cognitive functions, including cognitive control, emotion recognition, associative memory and cooperation [8–11]. However, the impact of these relatively brief laboratory taVNS experiments has been limited, for lack of evidence of a plausible mechanism of action. While animal studies have found that invasive VNS increases the firing rate of locus coeruleus neurons and increases extracellular norepinephrine levels [12–14], taVNS studies have generally found no or very small stimulation effects on non-invasive physiological markers of noradrenergic activity [15–18]—at least in studies using the relatively long on/off stimulation cycles of the taVNS device most often used in laboratory experiments (NEMOS®, Cerbomed GmbH, Erlangen, Germany). Positive findings of laboratory taVNS experiments with long on/off cycles or continuous stimulation, especially those using relatively low stimulation intensities (e.g., 0.5 mA) therefore warrant scrutiny, for example using direct replication studies.

Here we report a direct replication of a taVNS study [16] that has attracted a lot of interest, in particular because of its focus on gut-brain interactions, which have a major influence on motivational states. Previous evidence has shown that, as an important component of the autonomic nervous system, the vagus nerve is involved in allostasis, the regulation of motivation and energy metabolism [17,18]. Activation of vagal pathways can modulate reward-seeking behaviors [19], and control food intake through negative feedback signals routed via the NTS [20]. Thus, the vagus nerve may serve as a link between peripheral metabolic signals and central functions involved in goal-directed, allostatic behavior. To test the role of the vagus nerve in regulating motivation, Neuser and colleagues [1] examined the effect of taVNS on the motivation to work for rewards.

The participants in Neuser et al.’s study [1] carried out an effort allocation task in which they were asked, on each trial, to work for a primary (food) or secondary (money) reward by vigorously pressing a button. The reward magnitude and effort required to achieve the reward (i.e., task difficulty) varied across trials. After the effort phase of each trial, participants were asked to rate their wanting of the reward at stake and their exertion level (i.e., the perceived cost of their action). The 30-sec on phase of taVNS or sham stimulation was aligned with the 30-sec effort phase of the trial, followed by an off phase during the subjective ratings. Neuser et al. [1] found that taVNS boosted the invigoration of effort—that is, how quickly a participant energized effortful behaviour on each trial by ramping up the frequency of button presses. In contrast, taVNS did not affect the overall effort produced on each trial (i.e., average frequency of button presses).

The critical question in our study was whether we could replicate Neuser’s et al. [1] key finding that taVNS enhances the invigoration of effort. To ensure that we replicated the orginal study as closely as possible, we ran the task and analyzed the data using protocols and scripts kindly provided by Neuser and colleagues [1]. An important difference between the studies is that unlike Neuser et al. [1] we only tested participants with stimulation of the *left* cymba concha. In the past, the large majority of laboratory taVNS studies have only applied stimulation to the left cymba concha, because of cardiac safety concerns. Although these concerns have recently been challenged [20,21], our ethics committee did not allow us to administer taVNS to the right cymba concha. In this context it is important to note that the effects of left-sided and right-sided stimulation on invigoration reported by Neuser and colleagues [1] were essentially equal in size.

Before the start of the study, we decided to stop data collection when the evidence for or against the key hypothesis would reach a Bayes factor of 8 or 0.125 (i.e., 1/8), with a maximum of 40 included participants—Neuser and colleagues [1] tested 41 participants with left-sided stimulation. In accordance with these criteria, we stopped data collection after including 40 participants, and reaching a Bayes factor of 0.19, reflecting *substantial* evidence for the null hypothesis [22].

## Methods

### Participants

A total of 42 participants took part in the study, which consisted of two sessions scheduled between 2 and 8 days from each other at approximately the same time of day. In one session, participants received stimulation of their left cymba conchae (taVNS condition), in the other session, their left ear lobe was stimulated (sham condition). Two participants were excluded because they missed the second session or failed to respond on many trials. The resulting final sample consisted of 40 participants (32 women; mean age *M* = 21.30, range 18 to 29).

All participants were right-handed and used to eating breakfast between 6 and 10 am. Exclusion criteria were: having participated in any taVNS experiment in the previous month, history of cardiac conditions, neurological or psychiatric disorders, skin disorders, current use of psychoactive medication or drugs, and active implants (e.g., cochlear implant, pacemaker). Participants were asked to not consume any food or alcohol within the 12 hours before the start of the experiment. The study was approved by the ethics committee of the Institute of Psychology at Leiden University.

### taVNS stimulation

A NEMOS® stimulation device was used to stimulate the left auricular branch of the vagus nerve. Two titanium electrodes placed either at the left cymba conchae (taVNS) or left earlobe (sham) transmitted electrical current with biphasic impulse frequency of 25 Hz and a stimulation duration of 30 s [23]. This device has previously been used in both clinical settings [24,25] and fundamental research [1,23,26]. The default setting of the device follows a 30-s on phase followed by a 30-s off phase. To replicate Neuser’s et al. [1] procedure during the effort task, the experimenter aligned the onset of the stimulation to the start of each trial, which meant shortening the 30 s off phase.

Before participants performed the task, we carried out a work-up procedure to achieve a stimulation intensity that corresponded to a ‘mild pricking’ [1,26]. Participants were repeatedly asked to rate the received intensity of the stimulation on a 10-point visual analogue scale (“How strong do you perceive pain induced by the stimulation?” ranging from 0, “no sensation”, to 10, “strongest sensation imaginable”). Stimulation started at an amplitude of 0.1 mA and was increased in steps of 0.1-0.2 mA until a rating of 4, corresponding to a ‘mild pricking’ sensation, was reached (*M*_taVNS_ = 1.36 mA; range = 0.3 to 2.8 mA; *M*_sham_ = 1.84 mA; range = 0.6 to 4.4 mA). Participants then proceeded with the effort allocation task.

### Effort allocation task

Task scripts for administering the effort allocation task were provided by Neuser and colleagues [1] and were run using the Psychophysics Toolbox v3 [27,28] in MATLAB R2018b. During the task, participants were required to exert effort by repeatedly pressing a button with their right index finger on an Xbox 360 controller joystick (Microsoft Corporation, Redmond, WA), to obtain either monetary or food rewards. Like Neuser et al. [1], we measured the frequency of button presses as an indicator of physical effort. At the beginning of each trial, participants were presented with the reward that could be obtained on that trial (**Figure. 1**). This could be either food, represented by a picture of one or more cookies, or money, represented by a picture of one or more coins. The number of coins and cookies could be either one or several, indicating the magnitude of the reward on a given trial. Only one item corresponded to 1 point per second, whereas several items corresponded to 10 points per second.

**Figure 1.**
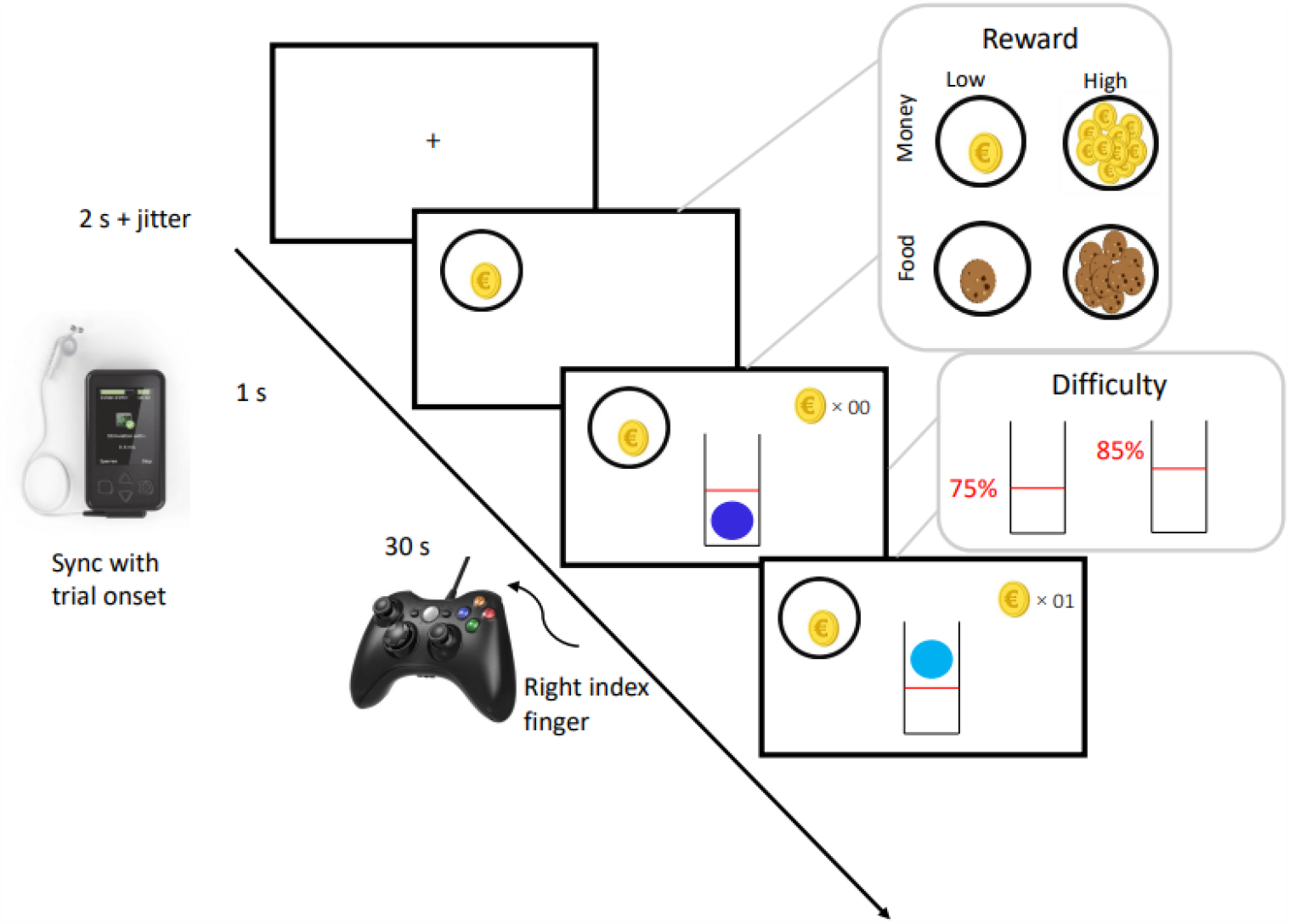
Schematic summary of the effort allocation task [1].

After 1 s, a blue ball inserted in a tube was shown on the screen. Participants were instructed to continually press a button with their right index finger to move the ball vertically up the tube and keep it above a red line. Depending on the level of difficulty for that trial, the height of the red line was adjusted. The difficulty was manipulated across trials (either 75% [low difficulty] or 85% [high difficulty] of the individual maximum frequency) with the order of difficulty counter-balanced across participants. At the start of each trial, the difficulty level was made clear by the height of the red bar. The longer the ball was kept above the line, the more reward points a participant earned on that trial. For every second the ball was above the red line, signalled by the ball changing colour from dark blue to light blue, reward points were added and the new total displayed in the top right corner of the screen.

Following the effort phase of a trial (30 s), two visual analogue scales were displayed on the screen. Participants were asked to rate their level of exertion, and their wanting of the reward on the preceding trial. The main experiment consisted of 48 trials interrupted by two short breaks of 15 s each. Before starting the task, participants were shown instructions, which emphasized that the task was too difficult to always keep the ball above the red line. Participants were encouraged to always hold the controller in the same way and to take breaks at their convenience to recover during trials, so that they could try to exceed the threshold again. At the end of the task, participants were shown the total amount of food and monetary rewards earned, with their conversions into euros and calories.

### Experimental procedure

The design of the current study was single-blind, within-subjects, randomized and counterbalanced across conditions. Both sessions started between 7:30 am and 10:10 am, as in Neuser et al. [1], and lasted around 1 h each. However, most of the participants were scheduled to start at either 8 am or 9:30 am. Participants were told they could drink water ad libitum during the experiment and that the earned points would be converted into calories and money at the end of the session.

Next, participants practised the effort allocation task for ∼5 minutes. During this training phase, the maximum frequency of button presses was estimated for each participant. A complete description of the practice phase can be found in Neuser et al. [1].

Next, we placed the taVNS device on the left ear of the participant, with the electrodes making contact with the cymba conchae (taVNS) or left earlobe (sham). To increase conductance of the electrodes, the participant was instructed to wipe their left ear with an alcohol pad. Moreover, two cotton rings were soaked with electrode contact fluid and applied on the taVNS device. Next, we assessed the stimulation intensity for each participant as described above. Then, participants completed the effort allocation task, which lasted ∼40 minutes. When asked at the end of each session which stimulation condition they were assigned to, participants were not able to guess better than chance (omitted guesses: 14, recorded guesses: 66, correct guesses: 32, accuracy: 48.5%).

At the end of each session, after completing the task, participants received their breakfast: a chocolate bar, milk, and cereals according to the number of calories earned. After the second session, participants also received their monetary compensation: a fixed amount of 22.50€ or course credits + the wins of both sessions. On average, participants won 562.52 kcal and €6.07 per session.

### Motivation indices and mixed-effects models of stimulation effects

To calculate the two motivational aspects, invigoration slope and effort maintenance, we used MATLAB R2021a and the original analysis scripts received from Neuser and colleagues [1], in which the behavioural data were divided into work segments and rest segments. Like Neuser et al. [1], we calculated the invigoration of effort by estimating the slope of the transition between the relative frequency of button presses during a rest segment and their initial plateau during the subsequent work segment (MATLAB findpeaks function). The maintenance of effort was calculated as the average frequency of button presses during a trial. In this way we estimated how much effort participants produced over time.

In accordance with Neuser et al. [1], invigoration slope and effort maintenance were moderately correlated, *r* = 0.28, 95% CI [0.25, 0.31]. The test-retest reliabilities of the two measures were high; the correlations across participants between session 1 and session 2 were 0.85 for invigoration and 0.92 for effort maintenance. To assess the effects of taVNS on invigoration and effort maintenance, single-trial estimates of the two variables were entered into two separate univariate mixed-effects models. Invigoration slope and effort maintenance were predicted as outcomes using the following dummy-coded predictors: stimulation (sham, taVNS), reward type (food, money), reward magnitude (low, high), difficulty (easy, hard), the interaction between reward magnitude and difficulty, and the interactions between stimulation and all other predictors. To account for the order of stimulation, we included stimulation order (taVNS first, sham first) at the participant level. To account for interindividual variance, we included intercepts and slopes as random effects. To examine the correlation between invigoration slope and effort maintenance with subjective ratings, we used mixed-effects models to predict either invigoration or effort maintenance as the outcome with wanting and exertion ratings as the predictors.

### Statistical threshold and software

Since the current study was an attempt at direct replication, our statistical thresholds and scripts for pre-processing data were identical to those used by Neuser et al. [1]. We used a two-tailed *α* ≤ 0.05 for the analysis of our primary research question of whether taVNS modulates invigoration slope or effort maintenance across conditions (i.e., the main effect of stimulation). Mixed-effects analyses were conducted with HLM v7 [29] and ImerTest in R [30]. To determine the relative evidence for one hypothesis over the other (i.e., whether or not taVNS facilitates motivational aspects of goal-directed behaviour such as invigoration or effort maintenance), we calculated Bayes factors (BFs) using one-sided Bayesian *t*-tests based on order-corrected individual estimates of all stimulation effects (calculated using ordinary least squares). To do that, we used the default Cauchy prior *r* = 0.707 in JASP v0.9 [31]. Results were also plotted with R v3.4.0 (i.e., ggplot; [32]).

## Results

### Effects of task variables on invigoration and effort maintenance

We first tested whether we found the same effects of task conditions on invigoration slope and effort maintenance as found by Neuser et al. [1]. In line with Neuser et al. [1], participants were quicker to invigorate behavior when they were working for larger rewards, *b* = 4.32, *t*(38) = 3.1, *p* = .004, BF_10_ = 10.54 (**Figure. 2a; Supplementary Table 1**), while invigoration did not differ between food and money trials, *b* = 1.97, *t*(38) = 1.38, *p* = .17, BF_10_ = 0.43 (**Figure. 2c**). Unlike Neuser et al. [1], we did not find an effect of difficulty on invigoration, *b* = -1.69, *t*(38) = -1.26, *p* = .22, BF_10_ = 0.53: participants were not slower to invigorate when the difficulty level was higher. Lastly, we found no association between invigoration and wanting ratings, *t*(38) = 1.29, *p* = .23, fixed-effects estimate = 0.116, or exertion ratings, *t*(38) = 0.06, *p* = .95, fixed-effects estimate = 0.005. In contrast, Neuser et al. [1] found an association between invigoration and wanting ratings, with a fixed-effect estimate of 0.213 (**Figure. 2e)**.

**Figure 2.**
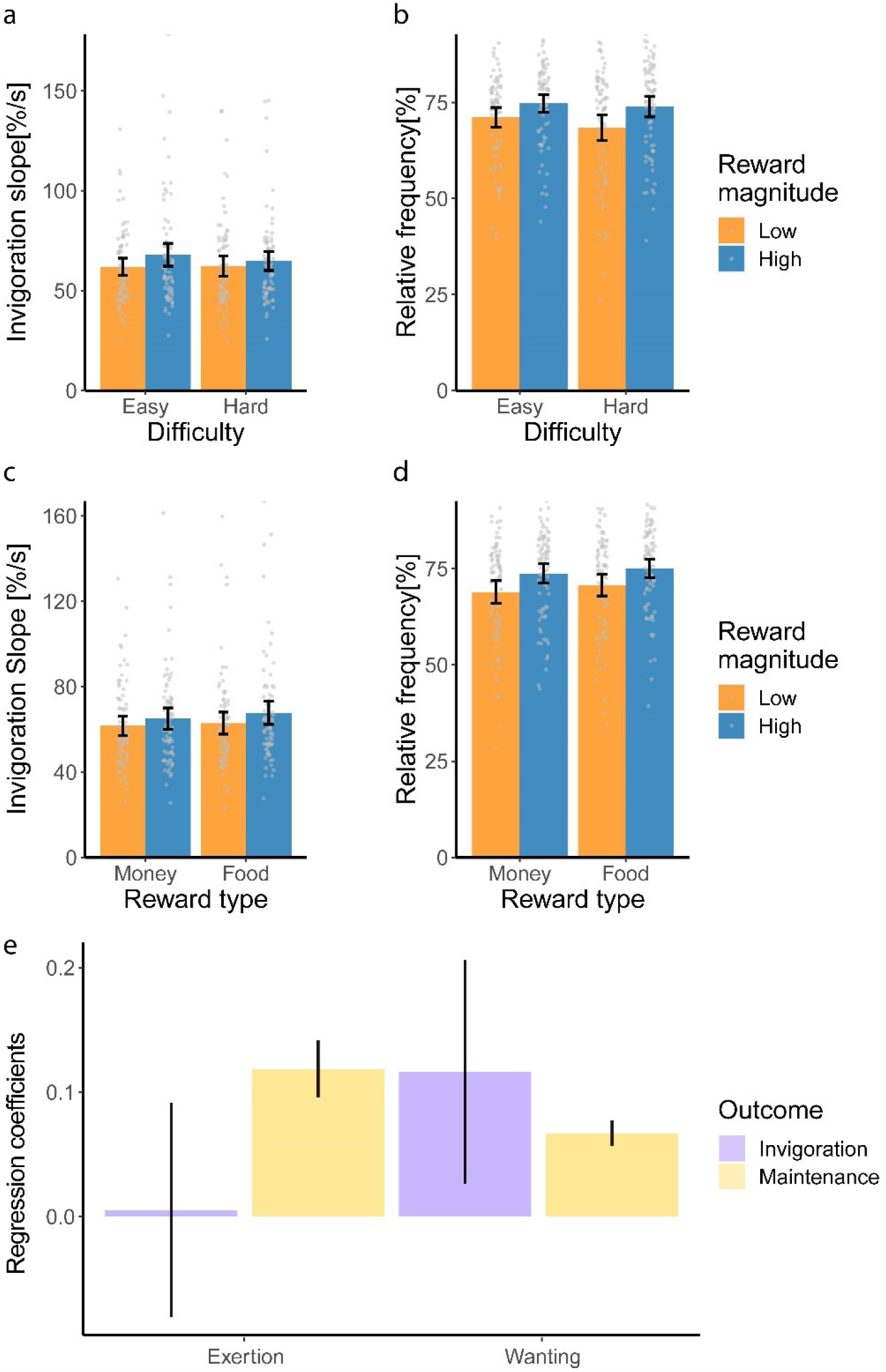
Invigoration is associated with reward magnitude while effort maintenance is linked with reward magnitude, trial difficulty and reward type. Grey dots indicate participant means per condition; error bars refer to 95% confidence intervals at the trial level (a-d). Bars indicate the fitted coefficients from the mixed models and error bars indicate the standard error of the mean (e). %/s = button press rate.

In line with Neuser et al. [1], participants showed higher effort maintenance when more reward was at stake, *b* = 4.4, *t*(38) = 4.01, *p* < .001, BF_10_ = 84.34, while their effort dropped significantly when reward was more difficult to obtain, *b* = -1.83, *t*(38) = -2.45, *p* = .019, BF_10_ = 1.79 (**Figure. 2b; Supplementary Table 2**). Furthermore, participants exerted more effort to obtain large rewards when the difficulty level was high, indicated by a reward magnitude × difficulty interaction, *b* = 0.95, *t*(38) = 2.65, *p* = .012, BF_10_ = 2.96 (**Figure. 2b**). Unlike Neuser et al. [1], we found higher effort maintenance when food as opposed to money was the reward at stake, *b* = 1.46, *t*(38) = 2.84, *p* = .007, BF_10_ = 5.88 (**Figure. 2d**), suggesting that food had a higher incentive value. In line with Neuser et al. [1], effort maintenance was associated with ratings of wanting, *t*(38) = 6.69, *p* < .001, fixed-effects estimate = 0.067, and exertion, *t*(38) = 5.17, *p* < .001, fixed-effects estimate = 0.119 (**Figure. 2e**).

### Effects of taVNS on invigoration and effort maintenance

Having replicated most of the behavioural effects reported by Neuser et al. [1], we next wanted to address the critical question of whether taVNS boosted invigoration. We found no effect of stimulation (taVNS vs. sham) on invigoration, *b* = 0.92, 95% CI [-1.04, 2.88], *t*(38) = 0.5, *p* = .62 (**Figure. 3a**). Importantly, a Bayesian analysis revealed substantial evidence for the null hypothesis, BF_10_ = 0.19. We also did not find the anticipated interaction between stimulation and reward type, *b* = -1.37, *t*(38) = -0.78, *p* = 0.44, BF_10_ = 0.23—a food-specific effect of taVNS on invigoration—which in Neuser et al. [1] was fully driven by left-sided stimulation. The impact of taVNS on effort maintenance was not influenced by any task factors (*ps* > 0.08, all BF_10_ < 0.99; **Figure. 4**).

**Figure 3.**
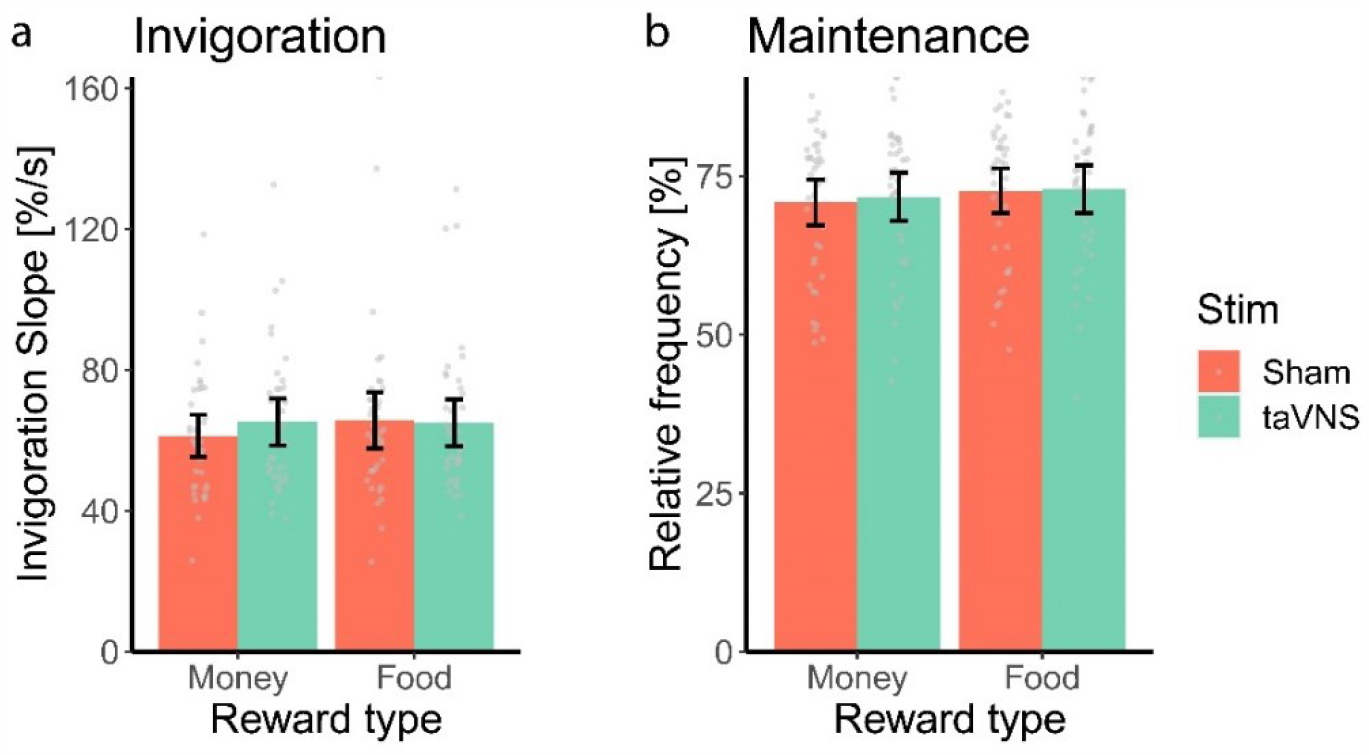
Effects of taVNS on the invigoration and maintenance of effort. Grey dots indicate participant means per condition; error bars refer to 95% confidence intervals at the trial level. %/s = button press rate.

**Figure 4.**
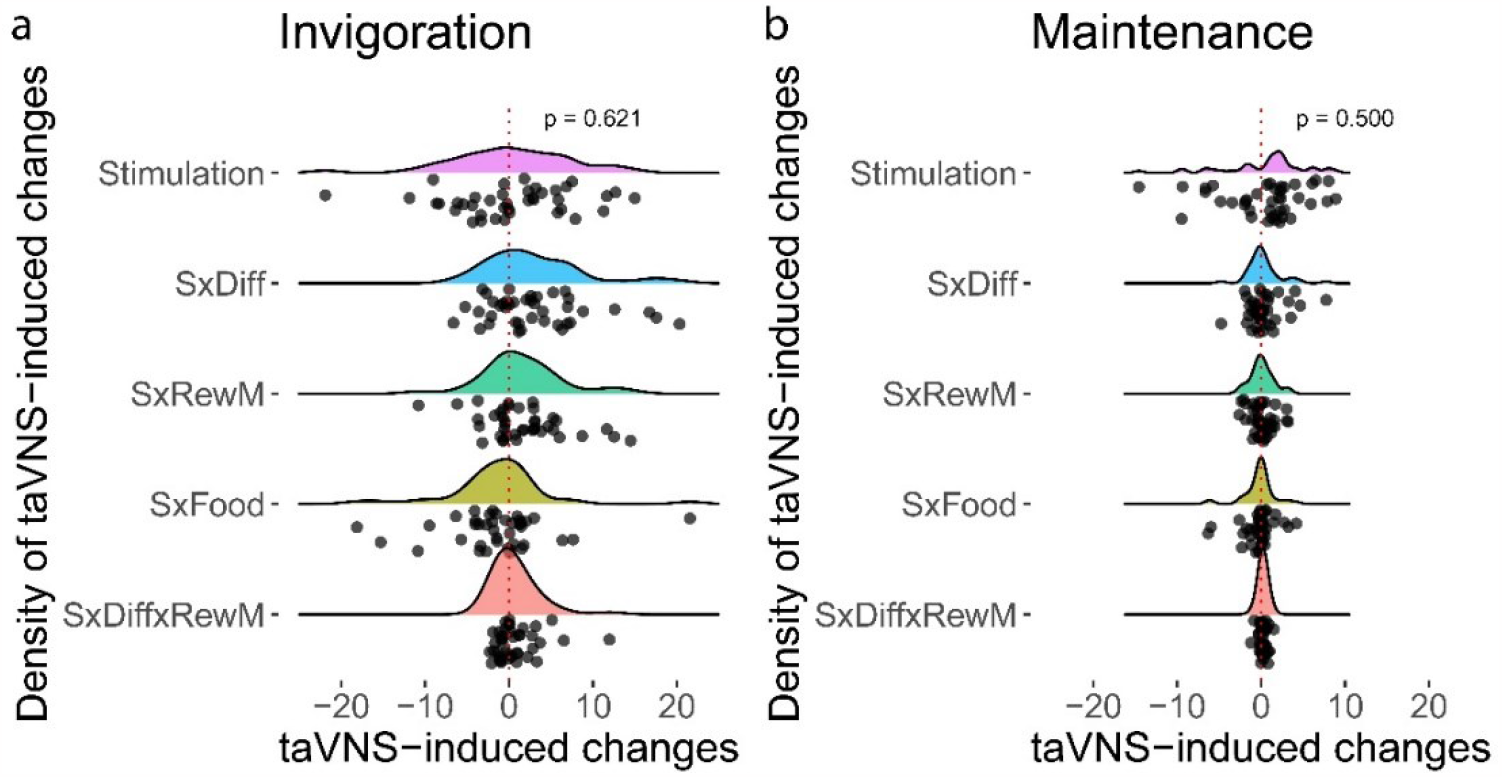
Stimulation effects induced by taVNS on invigoration and effort maintenance. Abbreviations: S = stimulation, Diff = difficulty, RewM = reward magnitude. P values refer to two-sided *t* contrasts of the mixed-effects models. Black points refer to the empirical Bayes estimates from the mixed-effects models.

Like in Neuser et al. [1], there was no effect of taVNS on effort maintenance, *b* = 0.48, *t*(38) = 0.68, *p* = 0.50; BF_10_ = 0.20; **Figure. 3b**) and the effect of taVNS on effort maintenance was not modulated by reward magnitude, difficulty or reward type (*ps* > 0.35, all BF_10_ < 0.25; **Figure. 4**).

## Discussion

Replicating scientific findings is crucial for scientific progress as it provides more robust knowledge, in our case about possible applications of tVNS in fundamental research and clinical practice. In the current study, we replicated the study of Neuser et al. [1], in which participants were required to exert effort by repeatedly pressing a button with their right index finger on a controller joystick to obtain either monetary or food rewards. Although, in line with Neuser et al. [1], we found strong effects of task variables on effort invigoration and maintenance, we were not able to replicate their key finding: taVNS did not increase the strength of invigoration, regardless of reward type, reward magnitude and trial difficulty. The data were five times more likely (BF_10_ = 0.19) under the null hypothesis than under the alternative hypothesis that taVNS modulates invigoration. Similar to Neuser et al. [1], we found no effect of taVNS on effort maintenance, although the relative strength of evidence for the null hypothesis was stronger in our data (BF_10_ = 0.20 compared to 0.51). Together, these findings cast doubt on the idea that active taVNS, compared to sham stimulation, can alter the effort or drive to work for rewards.

Importantly, for the most part, we were able to replicate the key task-related effects reported in Neuser et al. [1]. Notably, we found a robust effect of reward magnitude on invigoration and effort maintenance, suggesting that participants were more prone to invigorate their effort, and maintain this effort, when larger rewards were at stake. Furthermore, we replicated the effect of task difficulty on effort maintenance, showing that participants reduced the frequency of button presses when the trial was more difficult. We also replicated the null associations between subjective ratings of wanting and exertion and effort maintenance, but notably did not replicate the positive association between wanting and invigoration that Neuser and collagues [1] reported.

Our experimental procedure differed in a number of ways from that of Neuser et al. [1]. First, as discussed above, we were allowed to use only left-ear stimulation in our participants. We did not consider this a problem because Neuser and colleagues [1] found the effect of taVNS on invigoration to be essentially identical for left- and right-ear stimulation (interaction between taVNS/sham and side of stimulation: *p* = 0.947). Nevertheless, future research is necessary to determine if the reported positive effect of right-ear taVNS on invigoration is replicable.

Second, after stimulation was started, the participants in Neuser et al. [1] first carried out a 20-minute food-cue reactivity task [33] before they started the effort allocation task. This raises the possibility that we might also have found a positive effect of taVNS on invigoration if our participants had started the task after 20 minutes of stimulation. To address this possibility, we compared the effects of taVNS on invigoration during the first and the second 20 minutes of the effort allocation task by repeating the mixed-effects model analyses with an additional predictor (first vs. second half). These analyses yielded nonsignificant interactions between stimulation (taVNS/sham) and session half (first/second) for invigoration, *b* = 0.59, *t*(38) = 0.36, *p* = .79, and maintenance, *b* = 0.42, *t*(38) = 1.15, *p* = .26. These results suggest that 20 minutes of additional stimulation did not have substantial effects on the dependent variables.

Third, in our study, only 8 out of 40 (20%) participants were men, whereas in Neuser et al. [1] 34 out of 81 (42%) participants were men. Although this difference could potentially account for the discrepancy between our behavioral results, the limited available evidence suggests that female animals [22] and human participants [35] show larger effects of VNS than males, which is inconsistent with our null finding in a group of mainly female participants. ‘Invigoration’ and ‘vigor’ are well-developed concepts that generally refer to the (inverse) latency to initiate and/or complete a response, and have long been known to depend on dopaminergic activity [36]. It remains to be determined to what extent the invigoration slope in the effort allocation task used here, which does not reflect the latency of a discrete response, corresponds with these concepts. The moderate correlation with effort maintenance scores (*r* = 0.28), the high test-retest reliability (*r* = 0.85), and the significant effects of task variables suggest that invigoration slope is a meaningful motivational measure. Nonetheless, future research using more common measures of invigoration is needed to further assess the effects of invasive and non-invasive VNS on the motivation to work for monetary and food rewards.

Several recent taVNS studies have not managed to replicate the effects of invasive VNS on psychophysiological measures such as pupil size and P300 amplitude, reporting null effects instead [15–18,34]. In the present work and another replication attempt [35], we find that even results of high-profile taVNS studies can be difficult to replicate. Parametric exploration of tVNS stimulation parameters [36,37] and demanding authors to report internal replications may help this nascent field of inquiry to overcome these challenges.

## Data and code availability statement

Analysis code can be found here: https://github.com/bethlloyd/neuser_replication. Raw behavioral data will be made available upon publication.

## Acknowledgement

We would like to thank Anisha Koeldiep for her help with setting up the study. We would also like to thank the authors of the original paper [1] for their support in the replication of the study. Nils kindly provided us task and preprocessing scripts and gave us helpful advice on our analyses.

## Contributions

Conceptualization: S.N., B.L.; Data collection and curation: F.L.; Analysis: B.L.; Visualization: B.L.; Writing - original draft: F.L.; Writing - review and editing: S.N., B.L.; Supervision: S.N.

## Supplementary materials

**Supplementary Table 1.** Model output predicting invigoration

| Fixed effect | Coefficient | Standard error | t-ratio | Approx. d.f. | p-value |
| --- | --- | --- | --- | --- | --- |
| For INTRCPT1, $\pi_0$ | | | | | |
| INTRCPT2, $\beta_{00}$ | 61.517064 | 3.260963 | 18.865 | 38 | <0.001 |
| ORDER, $\beta_{01}$ | 0.552992 | 6.530231 | 0.085 | 38 | 0.933 |
| For CDIFF slope, $\pi_1$ | | | | | |
| INTRCPT2, $\beta_{10}$ | -1.688674 | 1.339186 | -1.261 | 38 | 0.215 |
| ORDER, $\beta_{11}$ | -2.020982 | 2.681834 | -0.754 | 38 | 0.456 |
| For FOOD slope, $\pi_2$ | | | | | |
| INTRCPT2, $\beta_{20}$ | 1.967329 | 1.421355 | 1.384 | 38 | 0.174 |
| ORDER, $\beta_{21}$ | 1.543758 | 2.846371 | 0.542 | 38 | 0.591 |
| For REWM slope, $\pi_3$ | | | | | |
| INTRCPT2, $\beta_{30}$ | 4.316805 | 1.390941 | 3.104 | 38 | 0.004 |
| ORDER, $\beta_{31}$ | 7.038127 | 2.785471 | 2.527 | 38 | 0.016 |
| For STIMCOND slope, $\pi_4$ | | | | | |
| INTRCPT2, $\beta_{40}$ | 0.924262 | 1.856471 | 0.498 | 38 | 0.621 |
| ORDER, $\beta_{41}$ | -8.091557 | 3.717671 | -2.177 | 38 | 0.036 |
| For I_DRM slope, $\pi_5$ | | | | | |
| INTRCPT2, $\beta_{50}$ | -1.416685 | 1.607062 | -0.882 | 38 | 0.384 |
| ORDER, $\beta_{51}$ | 1.215436 | 3.218239 | 0.378 | 38 | 0.708 |
| For I_SDIFF slope, $\pi_6$ | | | | | |
| INTRCPT2, $\beta_{60}$ | 2.817264 | 1.581673 | 1.781 | 38 | 0.083 |
| ORDER, $\beta_{61}$ | -4.553411 | 3.167400 | -1.438 | 38 | 0.159 |
| For I_SREWM slope, $\pi_7$ | | | | | |
| INTRCPT2, $\beta_{70}$ | 1.504684 | 1.574786 | 0.955 | 38 | 0.345 |
| ORDER, $\beta_{71}$ | 3.397860 | 3.153609 | 1.077 | 38 | 0.288 |
| For I_SFOOD slope, $\pi_8$ | | | | | |
| INTRCPT2, $\beta_{80}$ | -1.372511 | 1.760480 | -0.780 | 38 | 0.440 |
| ORDER, $\beta_{81}$ | -1.371749 | 3.525452 | -0.389 | 38 | 0.699 |
| For I_SDRM slope, $\pi_9$ | | | | | |
| INTRCPT2, $\beta_{90}$ | 0.817934 | 1.350391 | 0.606 | 38 | 0.548 |
| ORDER, $\beta_{91}$ | -1.963367 | 2.704271 | -0.726 | 38 | 0.472 |

**Supplementary Table 2.**
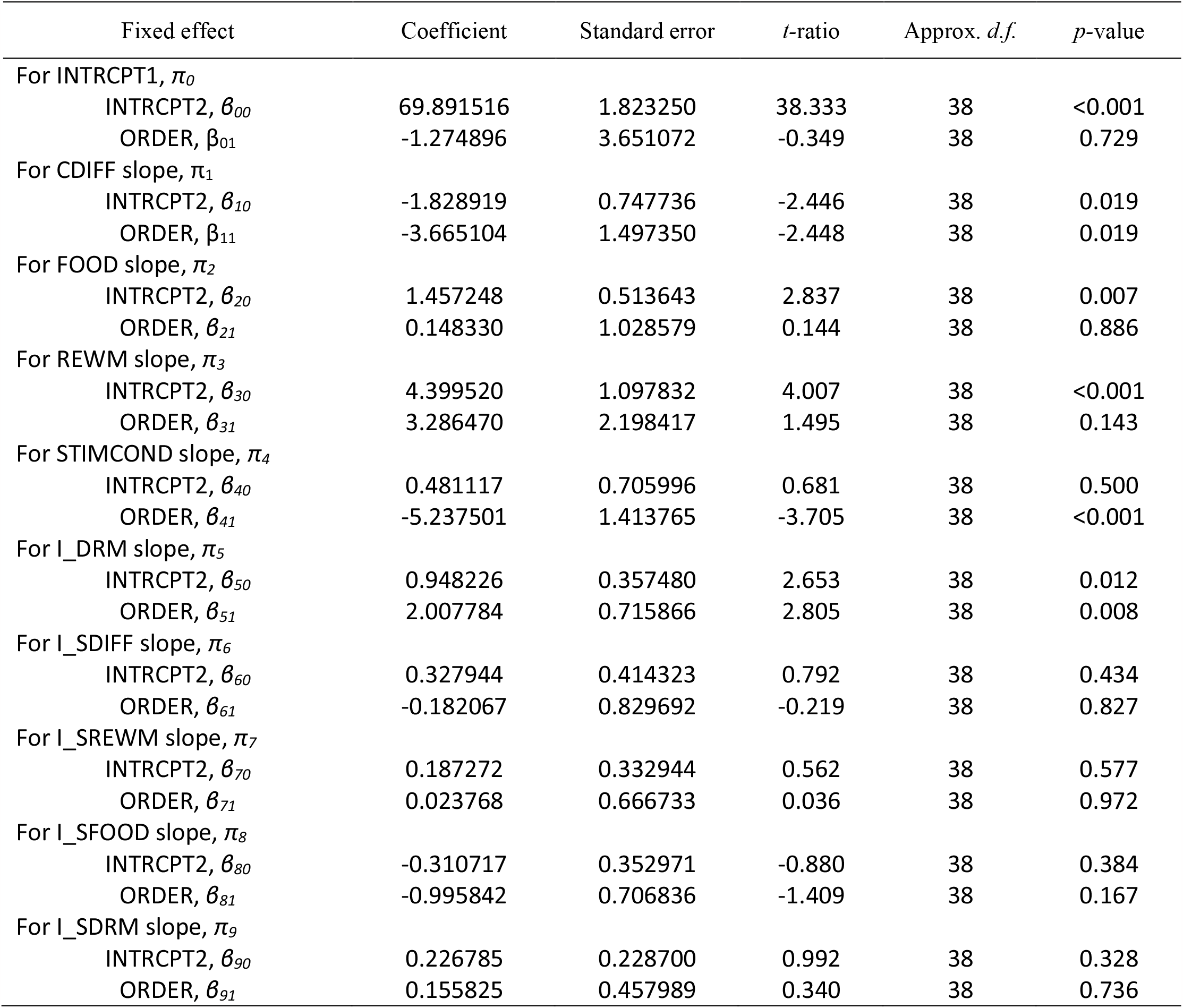
Model output predicting effort maintenance

